# Structural rationalization of IPMK inhibitor potency

**DOI:** 10.1101/2025.08.16.670669

**Authors:** Huanchen Wang, Stephen B. Shears, Raymond D. Blind

**Affiliations:** Molecular and Cellular Biology Laboratory, National Institute of Environmental Health Sciences, NIH, Research Triangle Park, NC 27709, USA; Department of Medicine, Division of Diabetes, Endocrinology & Metabolism, Vanderbilt University Medical Center, Nashville, TN 37232, USA; Departments of Biochemistry & Pharmacology, Vanderbilt University School of Medicine, Nashville, TN 37232, USA

**Author notes:** Corresponding Author Information*: Huanchen Wang., Ray Blind.

## Abstract

Inositol polyphosphate multikinase (IPMK) is a kinase linked to several cancers, recent development of a large panel of ATP-competitive inhibitors has reinvigorated enthusiasm for targeting IPMK. However, the structural basis for how these inhibitors achieve high potency is unknown. Here, we report 14 novel co-crystal structures (1.7Å - 2.0Å resolution) of human IPMK kinase domain with these inhibitors. We also apply a radiolabeled assay and isothermal titration calorimetry that permit high-confidence IC_50_ and K_D_ value determinations. The structures reveal a pocket in the ATP-binding site engaged by the most potent inhibitors. Two ordered waters also participate in hydrogen-bonding networks associated with the most potent inhibitors. In addition to providing the molecular basis for observed increases in potency and selectivity, the data presented here provide a toolbelt of 14 novel inhibitor-bound structures of human IPMK that can serve as a reference for all future IPMK structure-based inhibitor development efforts.

## INTRODUCTION

Inositol polyphosphate multikinase (IPMK) is a ubiquitously expressed kinase that has been linked to several cancers^1–10^, and loss of IPMK kinase activity in cells decreases cell growth and proliferation in several human cell lines. The recent development of a series of IPMK kinase inhibitors by another group has shown these inhibitors have demonstrated efficacy in decreasing cellular growth of human U251-MG glioblastoma cells and altering gene expression patterns in these living human cancer cells. Although this series of compounds are ATP-competitive, the structural details showing how these compounds interact with IPMK at the ATP binding site remain unclear. Atomic resolution details will be required for structure-based improvements on these compounds. That these compounds have demonstrated efficacy at inhibiting cancer cell growth suggests structure-based optimization may help translation as cancer therapies.

The first crystal structure of any IPMK orthologue published was that of the yeast ipk2^11^, followed by the plant *Arabidopsis thaliana* orthologue^12^. These structures established IPMK has the typical kinase fold, containing an N-terminal lobe and C-terminal lobe with the ATP binding site at the interface between these two lobes. The structure of the plant IPMK showed a larger IP-binding loop, suggesting why the plant orthologue has no activity on the phospholipid PI(4,5)P2^7^, yet retains inositol phosphate kinase activity^12^. However, crystallographic analysis of the yeast and plant enzymes, lacking bound inositol phosphates, does not provide a structural rational for activities of human IPMK on PI(4,5)P2. X-ray crystal structures of the human IPMK kinase domain were solved in close succession by two independent labs, both in the apo form^13^ and in complex with nucleotide^14^, and with the kinase substrates inositol phosphate and the phosphoinositide lipid PI(4,5)P2^14^. These structures, when associated with extensive kinetic analyses of IPMK activity, revealed several interesting aspects of IPMK catalysis and regulation. However, still no small molecule inhibitor-bound structures of IPMK had been solved to that point.

X-ray structural analyses of human IPMK bound to several different ATP-competitive flavonoids, which are general inhibitors of many diverse kinases, revealed hydrophobic and polar interactions between the flavonoids and particular amino acid side chains in IPMK^15^. These studies also suggested that ordered water molecules in the ATP-binding site might be important for flavonoid interaction with IPMK, and informed potential pharmacophore properties in the development of inositol-phosphate kinase inhibitors. However, flavonoids are relatively non-specific protein and phospholipid kinase inhibitors, so structure-based improvements would likely require not only improvements to flavonoid interactions with human IPMK, but also chemical modifications that would discourage interaction with other kinases, to have the best chance at clinical utility. Thus it would be useful if novel chemical compounds could be developed that were specific to IPMK.

In the process of identifying inhibitors of IP6K1, a close structural relative to IPMK within the inositol-kinase superfamily^16,17^, we found that some compounds synthesized as inhibitors of IP6K1 were effective IPMK inhibitors. Those compounds were recently optimized using medicinal chemistry, producing several generations of IPMK inhibitors with pharmacokinetic properties that suggest potential for clinical translation^18^. However, the lack of any X-ray crystal structural information of these compounds dramatically limits the potential for structure-based rational improvements to these compounds, potentially impeding progress to the clinic.

Here, we solved the X-ray crystal structures of 14 of this new generation of compounds complexed with the kinase-domain of human IPMK. We also apply a highly quantitative ^33^P-radiolabel-based binding assay that permits accurate estimation of the IC_50_ for these compounds, as well as isothermal titration calorimetry to establish the K_D_ for the first generation compound binding to purified human IPMK^17^. The 14 crystal structures have revealed novel motifs within the IPMK inhibitor compounds that will encourage future structure-based optimization of these compounds. Together, the data presented here provide a definitive framework for the structure-based development of IPMK inhibitors.

## RESULTS

### Establishing IC50s for selective of IPMK inhibitors

Previous efforts developed kinase inhibitors of IP6K1 in collaboration with another group produced compound **1**^19^. In this study, we determined the IC_50_ values using a ^33^P-radioloabled HPLC-based assay (see methods) to overcome challenges in detecting the radiolabeled products of the kinase reactions (**Fig 1A**). Compound **1** IC_50_ values suggest more potency toward IPMK (IC_50_=12.8nM) than either of the related kinases IP3KA (IC_50_=8000nM) or IP6K1 (IC_50_=146nM) (**Fig 1B**), with compound 1 having approximately 10-fold selectivity for IPMK over IP6K1. Compound **1** directly bound IPMK with K_d_=11nM by isothermal titration calorimetry (**Fig 1C**), consistent with IC_50_ value. These data suggest that compound **1** preferentially binds recombinant human IPMK kinase domain, when compared to the structurally related human IP6K1 enzyme. However, without a structure, the details of the interaction with IPMK remain unclear.

**Figure 1:**
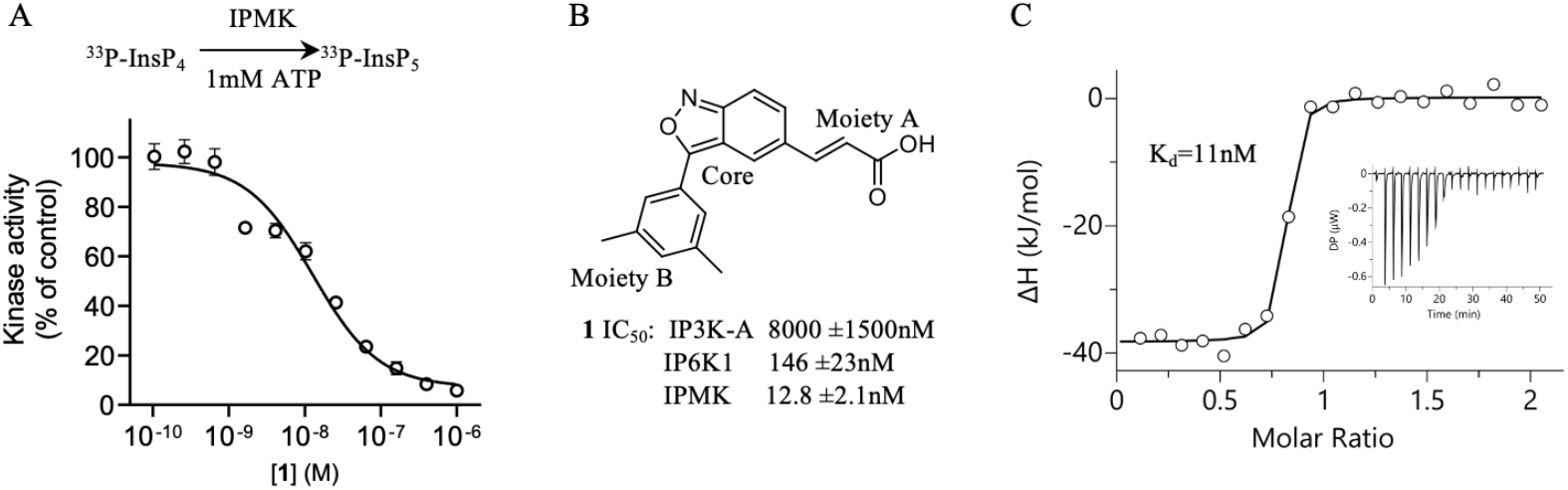
Compound 1 binds directly to IPMK with high affinity and effectively inhibits IPMK kinase activity. **A**. Top shows overview of ^33^P-radiolabeled HPLC assay used to determine IC_50_ values throughout this study. Lower plot of the IPMK kinase activity as a function of compound **1** concentration, determined by HPLC of substrate ^33^P-InsP_4_ conversion to product ^33^P-InsP_5_. **B**. Chemical structure of compound **1**, IC_30_ values for inhibition of in vitro kinase activity of purified IP3K-A, IP6K1 and IPMK, as indicated, error represents standard error from n=3 independent measurements. **C**. Isothermal Titration Calorimetry (ITC) plot of measured changed in enthalpy (ΔH) as a function of the molar ratio of compound 1 to IPMK, showing saturable binding of compound 1 to IPMK, inset shows a representative ITC isotherm, some error bars are obscured by the data symbols. *These data suggest compound 1 effectively and potently inhibits the kinase activity of purified human IPMK enzyme*.

### Co-crystal structure of compound 1 with human IPMK kinase domain

In order to understand how compound **1** interacts with IPMK for structure-based compound development, we solved the 1.95Å X-ray crystal structure of compound **1** in complex with human IPMK (**Fig 2A**, Supplemental **Table S1** for all crystallography statistics). As expected and established in other crystallographic studies of IPMK^15^, the nucleotide-binding site of human IPMK adopts the typical kinase N-lobe and C-lobe fold connected by a hinge loop (**Fig 2A-B**) and the nucleotide binding site (**Fig 2B**) occupied by an ambiguous electron density interpreted as compound **1** (**Fig 2C**). Compound **1** contains three chemical moieties, the benzisoxazole ring we call the core, with moieties A (acrylic acid) and B (dimethylphenyl) on either side (**Fig 2B**). The core of compound **1** formed four hydrogen bonds with the IPMK hinge region, including three hydrogen bonds with IPMK backbone residues and one hydrogen bond with the carboxylic group of Asp132 (**Fig 2C-E**). For comparison, the adenine group of ATP in the co-crystal structure with IPMK only makes 2 hydrogen bonds with the polypeptide backbone^14^. The overall fold of the IPMK kinase domain is similar to those structures already published in complex with various other small molecules and substrates.

**Figure 2.**
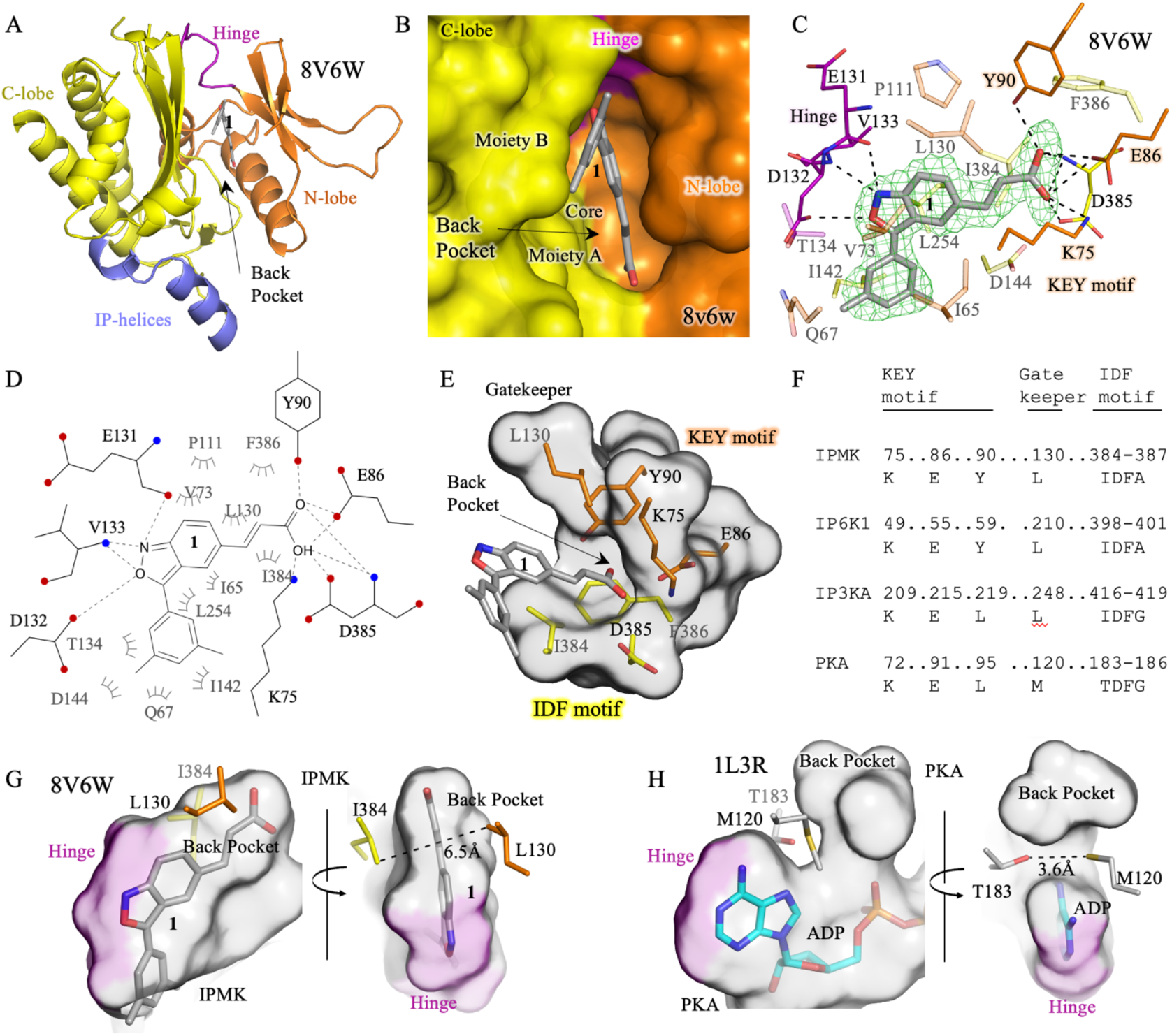
The 1.95Å crystal structure of compound 1 with IPMK (8V6W) reveals conserved residues in IP-kinases that mediate interaction with compound 1. **A**. Ribbon representation of the human IPMK co-crystal structure with compound **1**, various regions of IPMK labeled as indicated to provide frame of reference. **B**. Surface representation of IPMK-compound **1** co-crystal structure, with the N-terminal lobe colored orange and the C-terminal lobe colored yellow. **C**. Compound **1** ligand view showing indicated contacts in the co-crystal structure between compound 1 and indicated human IPMK residues, density throughout the figure represents the F_o_−F_c_ electron density omit map, generated by excluding ligands from the model, contoured at 3.0σ. **D**. Ligplot of identical region as in panel C. **E**. Protein-only surface representation of the same compound **1-**binding region in the human IPMK co-crystal structure. **F**. Sequence alignment of the KEY motif with corresponding amino acid numbering for indicated human kinases (IPMK, IP6K1, IP3K-A and PKA). **G**. Culled surface representation of compound **1** binding pocket, labeled as indicated. **H**. Culled surface representation of Protein Kinase A bound to ADP for comparison (1L3R), labeled as indicated with the hinge domain colored purple. *These data suggest compound 1 binds to IPMK via the KEY motif, which is a conserved motif in several IP-kinases*.

### Identification of anchored compound 1 binding and occupancy of a unique sub-pocket

The crystal structure of compound **1** with IPMK also revealed that moiety A penetrated a sub-pocket that is contoured by three hydrophobic IPMK residues (**Fig 2E**, Leu130, Iso384, and Phe386). The four polar amino acids in the IPMK back-pocket (Lys75, Glu86, Tyr90, and Asp385, **Fig 2C-E)** also formed a hydrogen bond network with moiety A of compound **1**. The residues Lys75, Glu 86, and Asp385 comprise the catalytic triad in IPMK, these residues are conserved in inositol phosphate kinases. ^13,20^. However, Tyr90 is unique to IPMK and IP6K, and is not present in IP3K or the related PKA protein kinase (**Fig 2F**). Thus, we named residues Lys75, Glu86 and Tyr90 the “KEY motif”, with this motif specific to inositol phosphate kinases. Together with the hinge region, IPMK anchored compound **1** with two hydrogen bond clusters, suggesting a two-point anchoring recognition mechanism. The core of compound **1** also forms several hydrophobic interactions with residues Ile65, Val73, Pro111 and Leu254 of IPMK, also observed in the complex of ATP with IPMK. In addition, Leu130, Ile384 and Phe386 are close enough to form van der Waals contacts with moiety A. For moiety B, Gln67, Thr134, Ile142, Asp144 and Leu254 are within van der Waals contact range. Thus, the co-crystal structure of compound **1** with human IPMK reveals multiple structure-based elements that can be used to improve the potency of compound **1** in future analyses.

### Crystal structures of first generation compounds 2-4 reveal the importance of ordered water molecules

Another group recently synthesized a series of several variants of compound **1** who have graciously shared these compounds. All compounds characterized in this study were synthesized and provided for our use by that group, which has been recently reported^18^. Here, we determined the potency of these compounds against IPMK and IP6K1 using the ^33^P-radiolabeled HPLC assay (**Fig 3A**). Compound **2** is more potent against both IPMK and IP6K1 than compound **1** (**Fig 3B**), however compound **3** had worse potency for IPMK (**Fig 3A-B**), while compound **4** had improved potency against IPMK (**Fig 3A**). To establish the structural determinants of these differences, we solved three new X-ray co-crystal structures of human IPMK complexed with compound **2** (1.75Å), compound **3** (1.70Å) and compound **4** (1.85Å) (**Fig 3C-F**). These high-resolution X-ray structures show a well-coordinated water labeled as water1. Compound **4** formed two hydrogen bonds with water1, while compound **2** and compound **3** coordinated water1 with only one hydrogen bond. The structures also suggest the acid groups in moiety A of compounds **2** and **3** are in proper position to have electrostatic repulsion with the acidic amino acid side chains of IPMK residues Glu86 and Asp385. The X-ray structure of IPMK with compound **4** suggested another water2 molecule with less well-defined density close to moiety A (**Fig 3E**) and revealed four hydrogen bonds between compound **4** and IPMK, consistent with compound **4** as the most potent inhibitor of these four compounds. Together, these data identified the IPMK residues most important to the improved inhibitory activity of compound **4**, and further support an important role of ordered water molecules in the binding mechanism for this series of highly potent compounds.

**Figure 3.**
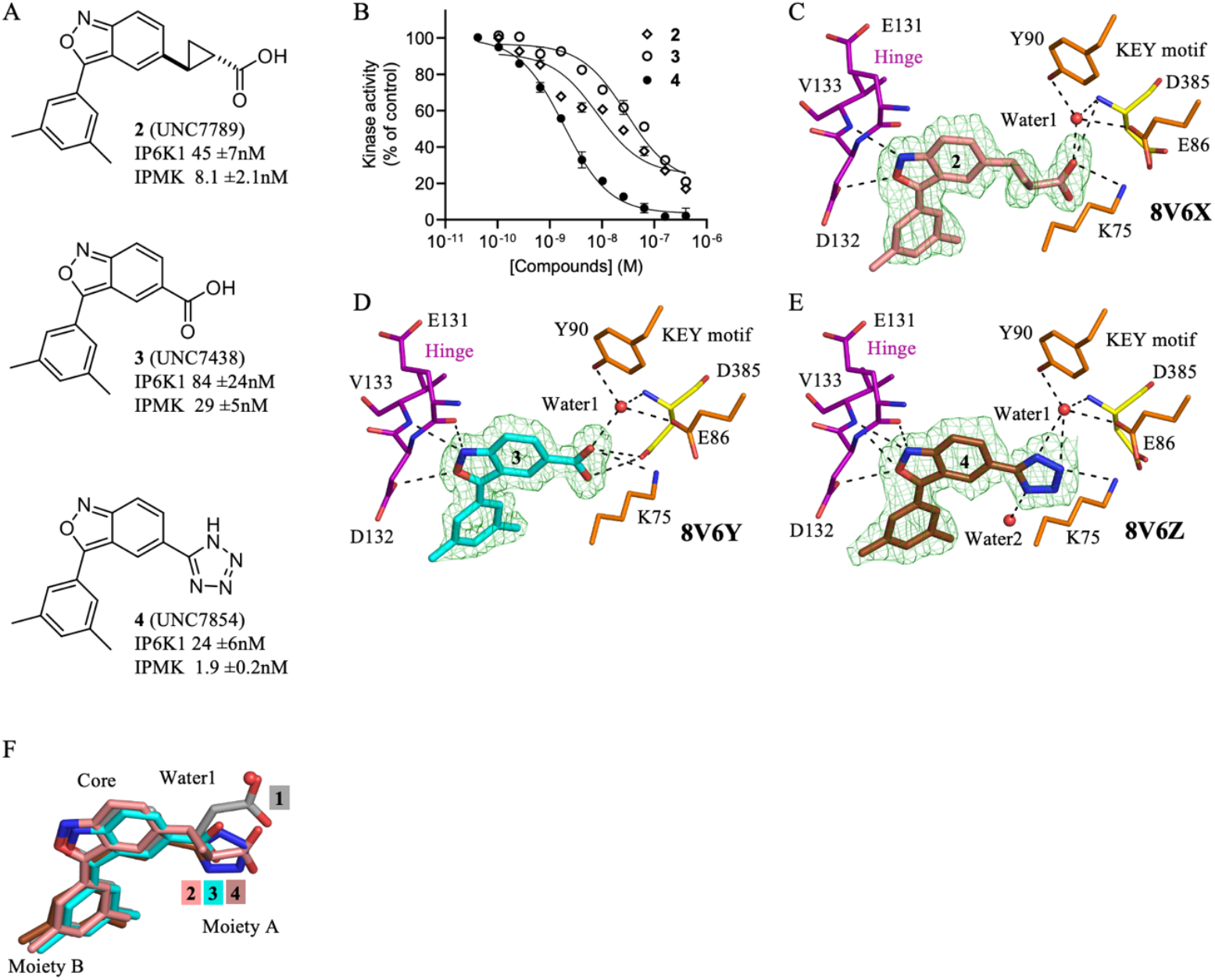
Modifications to Compound 1 show improved potency and efficacy. **A**. Chemical structures of **2, 3** and **4**, IC_50_ values for inhibiting ^33^P-radiolabeled HPLC in vitro kinase assay indicated, n=3, error represents standard error. **B**. Full curve fits of **2, 3** and **4** used to determine IC_50_ values in A, some error bars are obscured by the data symbols. **C**. Position of **2** in the 1.75Å co-crystal structure with human IPMK (8V6X), green mesh is electron density assigned to **2**, density throughout the figure represents the F_o_−F_c_ electron density omit map, generated by excluding ligands from the model, contoured at 3.0σ. **D**. Position of **3** in the independent 1.70Å co-crystal structure with human IPMK (8V6Y), green mesh is electron density assigned to **3. E**. Position of **4** in the 1.85Å co-crystal structure with human IPMK (8V6Z), green mesh is electron density assigned to **4. F**. Superposition of **1, 2, 3** and **4**, with labels indicating moiety A, moiety B and the core of the compounds. *These data suggest all compounds bind the ATP-site in IPMK, and moiety A exists in more diverse positions than the core or moiety B*.

### Second generation compounds 6-15 have a wide range of activity and selectivity, driven by pocket occupancy and ordered waters

The synthesis of second generation compounds 6-15 by another group were directed towards improving potency against IPMK and improving drug-like properties(18). We determined IC_50_ values for compounds 6-15 using the ^33^P-radiolabeled HPLC assay for IPMK and IP6K1 for comparison. Compound 6 had an IC_50_ for IPMK of 3.8nM, with ten-fold selectivity for IPMK over IP6K1 (Table 1). Compounds 7 and 8 had similar IC_50_ values and IPMK selectivity over IP6K1 when compared to compound **6**. Compound **9** had three-fold worse IC_50_ for IPMK but had increased selectivity over IP6K1. Compound **10** had similar selectivity over IP6K1 compared to **6**, while compound **11** had improved IC_50_ against IPMK (0.99nM) and fifty-fold selectivity over IP6K1. Compound **15** was similar in potency to **11**, compound **12** was equally potent and selective while compound **13** was three-fold less potent, and five-fold less selective for IPMK over IP6K1 compared to compound **4**. Compound **14** retained potent inhibitory activity against IPMK with an IC_50_ of 3nM, along with 52-fold selectivity over IP6K1. Together this wealth of compound inhibitory activity on IPMK provides baseline inhibitory data for all compounds in **Table 1**, however these data do not structurally rationalize the basis for the potency differences observed in the compounds.

**Table 1.** Modifications of Moiety B (R position) increase selectivity for IPMK over IP6K1.

| Compound | R | UNC# | IPMK<br>IC<sub>50</sub> (nM)<sup>a</sup> | IP6K1<br>IC<sub>50</sub> (nM)<sup>a</sup> | IP6K1/IPM<br>K |
| --- | --- | --- | --- | --- | --- |
| **1<sup>b</sup>** | N/A | UNC7437 | 12.8 ±2.1 | 146 ±23 | 11.4 |
| **2<sup>b</sup>** | N/A | UNC7789 | 8.1 ±2.1 | 45 ±7 | 5.6 |
| **3<sup>b</sup>** | N/A | UNC7438 | 29 ±5 | 84 ±24 | 3 |
| **4<sup>b</sup>** |  | UNC7854 | 1.9 ±0.2 | 24 ±6 | 13 |
| **6** |  | UNC7782 | 3.8 ±0.4 | 38 ±8 | 10 |
| **7** |  | UNC7855 | 2.5 ±0.3 | 23 ±4 | 9.2 |
| **8** |  | UNC7891 | 3.3 ±0.4 | 59 ±14 | 13 |
| **9** |  | UNC7890 | 11 ±2 | 1270 ±330 | 115 |
| **10** |  | UNC8787 | 3.5 ±0.4 | 35.8 ±5.8 | 10 |
| **11** |  | UNC9021 | 0.99 ±0.17 | 51.8 ±5.0 | 52 |
| **12** |  | UNC9120 | 3.4 ±0.5 | 29.9 ±6.0 | 8.8 |
| **13** |  | UNC9107 | 9.5 ±2.1 | 19.0 ±4.1 | 1.9 |
| **14** |  | UNC9750 | 2.8 ±0.6 | 155 ±23 | 52 |
| **15** |  | UNC9022 | 1.13 ±0.21 | 40.1 ±4.4 | 36 |

<sup>a</sup> Values are the mean of three or more independent assays with standard error, for Compound 1, n=5. For all other compounds, n=3. <sup>b</sup> Compounds 1-3 vary at the tetrazole shown (Moiety A, see Fig 3A for structures of Compounds 1-3), IC<sub>50</sub> data are provided here for comparison across all 14 compounds examined in this study.

### Crystal structures of second generation compounds reveal atomic-resolution details of IPMK inhibitor mechanisms

To address how these various modifications resulted in the observed changes to inhibitory activity, we determined the co-crystal structures of human IPMK with a total of 14 of these different IPMK inhibitors ranging in resolution of 1.75Å to 2.0Å (see **Supplemental Table S1** for crystallography statistics). These structures revealed specific details on the binding mechanism of each of these compounds, elucidating how they interact with IPMK and further highlighting the important role of ordered water molecules in compound binding. Crystal structures show that water1 persists across these compounds, except compound **1**. The structures of compound **6** (1.85Å, **Fig 4A-C**), compound **7** (1.70Å, **Fig 4D**) and compound **8** (2.0Å, **Fig 4E**) bound to IPMK all show a coordinated water2 molecule in the active site, similar to structures of compound **9** (1.90Å, **Fig 5A-D**) and compound **10** (1.90Å, **Fig 5E**). However, the structure with compound **11** showed water2 and a new water3 molecule in the active site, forming an additional hydrogen bond network with Arg182 (1.85Å, **Fig 5F**). These extra interactions, summarized in **Supplementary Table S2**, may account for the higher potency of compound **11**. In contrast, the structure with compound **15** (1.9Å, **Fig 6E**) still had water2 and water 3 ordered but did not show a detectable interaction with Arg182 (**Fig 6E**). The additional water3 molecule, together with water2, forms an internal hydrogen bond network between moieties A and B in both compound 11 (**Fig 5F**) and **15** (**Fig 6E**) structures. These water interactions may contribute to the particularly high potency of compounds **11** (**Fig 5F**) and **15** (**Fig 6B-E**). The co-crystal structure of compound **12** with IPMK (1.95Å, **Fig 6C**) confirmed **12** is accommodated in the binding site but did not gain any additional interactions with the IPMK protein when compared to compound **11** (**Fig 5F**) or compound **15** (**Fig 6E**). The structure of compound **13** with IPMK shows the meta-3,5-dichloro-pyridinyl group on moiety B of compound **13** makes two polar contacts with the backbone of Gly180 and Met181 (**Fig 6D**). The structure of compound **14** with IPMK (1.95Å, **Fig 5G**) shows water2 forming an internal hydrogen bond network similar to that seen with compound **11** (**Fig 5F**) and **15** (**Fig 6E**). Together, these 14 co-crystal structures of IPMK bound to these inhibitors have revealed the mechanism each compound uses to interact with IPMK, the importance of ordered waters in the active site of IPMK for binding inhibitors and opportunities for structure-based development of future IPMK kinase inhibitors.

**Figure 4.**
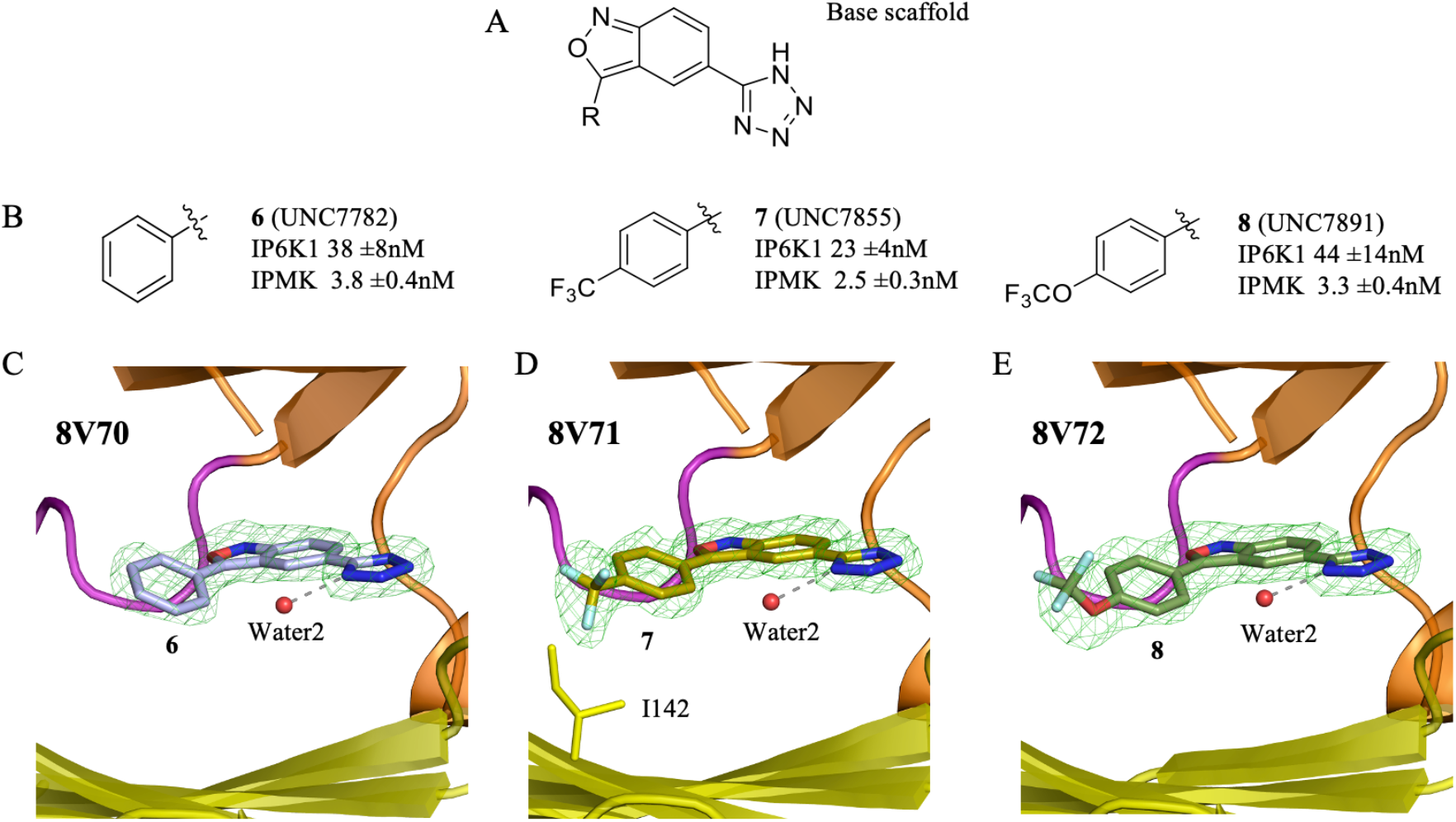
Crystal structures of IPMK with compounds 6, 7 and 8 show an ordered water2 in the active site. **A**. Chemical structure for the base scaffold. **B**. Chemical structures and IC_50_ values for IPMK *in vitro* kinase activity for compounds **6, 7** and **8**, n=3 error represents standard error. **C**. Position of compound **6** in the 1.85Å co-crystal structure with human IPMK (8V70), green mesh represents electron density assigned to compound **6**, red sphere represents ordered water2 molecule. **D**. Position of compound **7** in the 1.70Å co-crystal structure with human IPMK (8V71), green mesh represents electron density assigned to compound 7,1142 shown as stick representation, red sphere represents ordered water2 molecule, density throughout the figure represents the F_o_−F_c_ electron density omit map, generated by excluding ligands from the model, contoured at 3.0σ. **E**. Position of compound **8** in the 2.0Å co-crystal structure with human IPMK (8V72), green mesh represents electron density assigned to compound **8**, red sphere represents ordered water2 molecule. *These data suggest compounds **6, 7** and **8** all bind the active site of IPMK and highlight the role of ordered water2 molecule in compound binding*.

**Figure 5.**
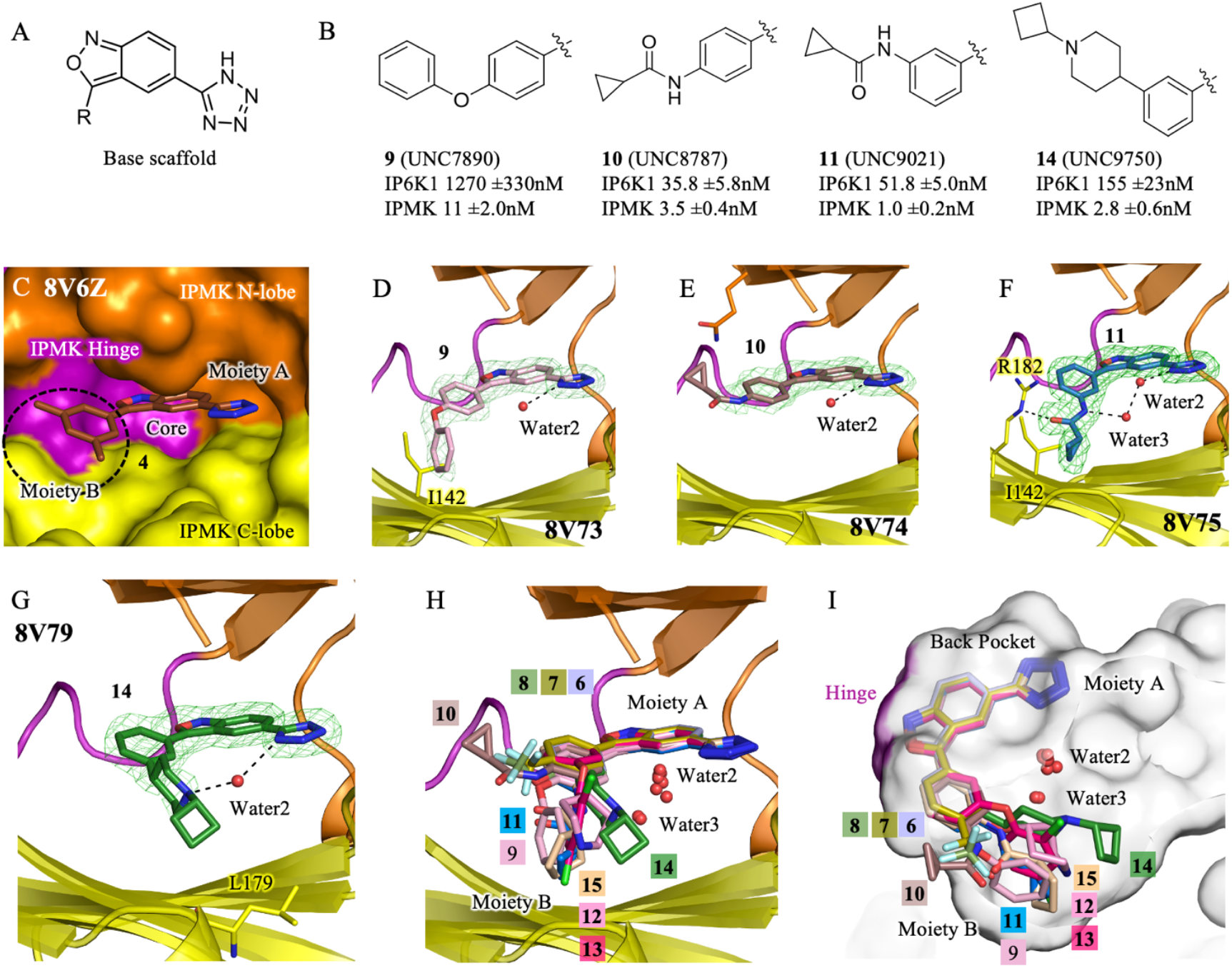
Crystallography reveals that further compound modifications have diverse positioning bound to IPMK. **A**. Chemical structure for the base scaffold. **B**. Second generation compounds **9**,**10**,**11** and **14**, with indicated IC_50_ values, n=3, error represents standard error. **C**. Surface model based on the 1.85Å X-ray crystal structure of IPMK and compound **4**, IPMK domains and compound **4** moieties as indicated. **D**. Position of 9 and water2 in the 1.90Å co-crystal structure with human IPMK, green mesh is electron density assigned to **9**, density throughout the figure represents the F_o_-F_c_ electron density omit map, generated by excluding ligands from the model, contoured at 3.0σ. **E**. Position of **10** in the 1.85Å co-crystal structure with human IPMK, green mesh is electron density assigned to **10. F**. Position of **11** in the 1.85Å co-crystal structure with human IPMK, green mesh is electron density assigned to **11. G**. Position of **14** in the 1.95Å co-crystal structure with human IPMK, green mesh is electron density assigned to **14. H**. Superposition of co-crystal structure position of compounds **6–15** (8V70, 8V71, 8V72, 8V73, 8V74, 8V75, 8V76, 8V77,8V78, 8V79), with water2 and water3 indicated, IPMK coloring identical as panel C. **I**. Culled surface representation of panel H, turned counterclockwise about the y-axis. *These data highlight the conformity in compound position at Moiety A, and the diversity of positions that occur at Moiety B*.

**Figure 6.**
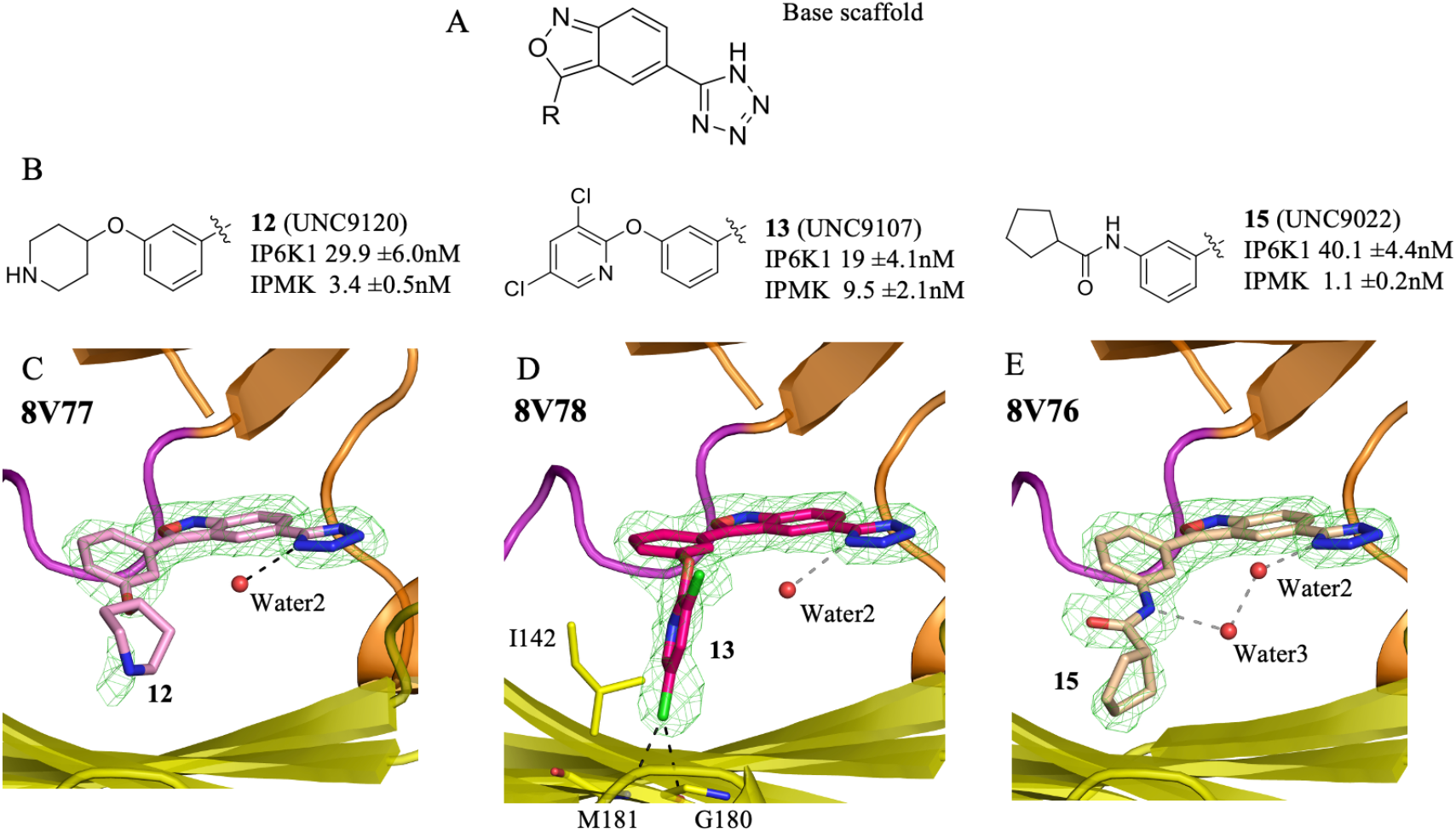
Crystal structures of IPMK with compounds 12, 13 and 15 show an ordered water2 in the active site. **A**. Chemical structure of the base scaffold. **B**. Chemical structures and IC_50_ values for inhibition of IPMK *in vitro* kinase activity for compounds **12**,**13** and **15**, n=3 error represents standard error. **C**. Position of compound **12** in the 1.95Å co-crystal structure with human IPMK (8V77), green mesh represents electron density assigned to compound **12**, red sphere represents ordered water2 molecule, density throughout the figure represents the F_o_−F_c_ electron density omit map, generated by excluding ligands from the model, contoured at 3.0σ. **D**. Position of compound **13** in the 1.96Å co-crystal structure with human IPMK (8V78), green mesh represents electron density assigned to compound **13**, red sphere represents ordered water2 molecule, 1142, Ml 81 and G180 shown as sticks. **E**. Position of compound **15** in the 1.90Å co-crystal structure with human IPMK (8V76), green mesh represents electron density assigned to compound **15**, red spheres represent ordered water2 and water3 molecules, as indicated. *These data suggest compounds* ***12, 13*** *and* ***15*** *all bind the active site of IPMK and highlight the role of ordered water3 in compound* ***15*** *binding*.

## DISCUSSION AND CONCLUSIONS

The mechanism of ligand recognition reported here involving a two-point anchoring mechanism with two polar contact clusters lays the groundwork for future inhibitor optimization targeting inositol phosphate kinases, including IPMK and IP6K. The core kinase domain of IPMK shares approximately 34% with IP6K1 and 24% sequence identity with IP3K-A, with high conservation observed in the ATP-binding site^14^. From the high-resolution structural data reported here, we highlight three broad observations that will be important for future structure-based drug design efforts directed against IPMK. First, moiety B of these compound scaffolds (**Fig 1B**) is open to the bulk phase (**Fig 5I**), and thus has more space to accommodate a wider variety of chemical groups and modifications. This available space imparts chemical flexibility at this position which may prove important in future iterations of IPMK inhibitor optimization. Second, the electron density around moiety A was consistently more well defined in the 14 crystal structures than the density around moiety B, suggesting interaction with IPMK has more flexibility at moiety B than at moiety A. Electron density of IPMK amino acids close to moiety B were as well defined as amino acids around moiety A, thus it is possible that modifications to moiety B designed to generate interaction with IPMK amino acids could improve potency, selectivity, and pharmaceutical properties. Third, chemical modifications at the para-position of the phenyl group in the compounds show less selectivity for IPMK, while modification at the meta-position shows increased selectivity for IPMK, as exemplified by the comparison between compound 10 (para-substitution) and compound 11 (meta-substitution), which showed a fivefold increase in selectivity. In all 14 crystal structures, the para-position of the phenyl group points to the bulk phase, while modification on meta-position could establish interactions with the residues from the C-terminal lobe of IPMK, where sequence differences with IP6K exist. Thus, it would be rational to attempt to increase compound selectivity for IPMK by introducing tailored modifications at both the para- and meta-positions.

We note that crystal structures of IP6K1 with the above compounds have not been determined, so we only formally describe interactions of the compounds with IPMK. However, applying computational prediction of human IP6K structures could guide future design of IP6K inhibitors as well. Indeed, we attempted to model compound **1** into the AlphaFold-predicted structures of IP3K-A and IP6K1 (**Supplementary Fig S1**). These models revealed a high degree of similarity in the binding sites, except that Tyr90 in IPMK is replaced by Leu219 in IP3K-A. In contrast, the hinge residue Asn212 in IP6K1 (corresponding to Asp132 in IPMK) may represent a key difference that helps discriminate between IP6K1 and IPMK. Most importantly, variations in the N-terminal residues, which are not visible in our crystal structures, may provide additional opportunities for achieving selectivity.

There are also some observations we have made that can be applicable more broadly to the kinase field. The hinge region, along with the C-spine, plays a significant role in inhibitor binding affinity in most protein kinases, as well as in inositol phosphate kinases^15^. The benzisoxazole ring core in this series established interactions, including polar contacts with hinge residues and hydrophobic interactions with C-spine residues. While these interactions serve as starting points for inhibitor design, compounds solely targeting the hinge region may lack selectivity, which can be achieved by targeting the back pocket. However, designing an inhibitor with back pocket binding presents additional challenges as has been previous noted^21,22^.

The back pocket, situated directly behind the adenine moiety, is also known as the hydrophobic pocket in protein kinases. In IPMK, the side chain of the gatekeeper Leu130 is relatively shorter than a methionine residue, allowing for more variation of moiety A in the back pocket^14^. The structures solved here show a polar contact cluster in this region, which together with essential polar contacts from the hinge region, suggest the two-point anchoring ligand binding mechanism proposed here to synergistically enhance potency of the IPMK inhibitors. Moiety B can expand in multiple directions without significant potency penalty. Water-mediated intramolecular interactions between moieties A and B may also affect pharmacokinetic properties.

Recent studies have demonstrated the set of IPMK inhibitors described here have efficacy in slowing human glioblastoma cancer cell growth, and regulate the expression of cancer-related gene sets^18^. However, the experimental crystallography and structural biology describing how these compounds bind IPMK has not been described, thus making any structure-based improvements to the compounds more difficult, and less confidence those modifications will actually improve compound potency or efficacy. The 14 crystal structures reported here provide the atomic-resolution details needed for these structure-based improvements to the inhibitors, making it more likely these compounds can be used clinically.

The previously published crystal structure of quercetin bound to IPMK showed a similar ordered water to that observed with the tetrazole group in this study^15^, indicating the potential importance of ordered water to IPMK ligand binding and selectivity. Importantly, several other kinase structures solved complexed with natural product flavonoids did not have a water molecule ordered in those structures^15^. In the co-crystal structures solved here, all compounds had ordered water molecules in the active site, Moreover, in compounds 11, 14, and 15, water2 and water3 form hydrogen bond networks that bridge the tetrazole group of moiety A with substituents on moiety B, thereby stabilizing the overall ligand conformation within the active site. These water-mediated intramolecular networks might help to lock the compounds into favorable binding poses even in the absence of direct contacts with IPMK residues, which likely contributes to their enhanced potency. The role of water in compound binding also highlights the power of X-ray crystallography in revealing compound binding mechanism.

Importantly, potency must be considered in tandem with selectivity. For example, compound 11 achieved the highest potency (0.99 nM) along with a marked improvement in selectivity over IP6K1, whereas compound 13 maintained moderate potency but showed substantially reduced selectivity. In contrast, compound 9 exhibited lower potency yet greater selectivity. These comparisons highlight that structural optimization of IPMK inhibitors cannot be guided by potency alone. Ideally, modifications that strengthen favorable interactions within the IPMK active site while minimizing contacts conserved in IP6K will achieve the optimal balance between potency and selectivity.

Recent work from another group that designed and synthesized this series of compounds also determined the pharmacokinetic parameters for compound **14**, revealing reasonable clearance, half-life and volume of distribution metrics after intraperitoneal injection in mice, suggesting that the pharmacokinetics and pharmacodynamics of this IPMK inhibitor has promise to translate to pre-clinical models of cancer in rodents, particularly glioblastomas^18^. Other studies have shown that human glioblastoma U251-MG cells decrease cellular growth and inositol phosphate metabolism in response to compound **14**, focusing clinical applications on glioblastomas might be more likely to successfully translate to pre-clinical rodent models^18^. However, significant data suggest that PTEN-negative cancers may respond generally to inhibitors of IPMK, as the kinase activity of IPMK is known to oppose the phosphatase activity of PTEN in the nucleus of U251-MG and HEK-293T cells^23^. Recent data also connect the kinase activity of IPMK specifically with the activity of HDAC3 in U251-MG cells, suggesting inhibitors of IPMK may synergize with HDAC inhibitors to bias the specificity of the response towards HDAC3^24–26^, although that hypothesis remains to be tested. Together, the data presented here reveal several fundamental structural principles of IPMK inhibitor design, new roles for ordered waters in IPMK inhibitor binding and provide atomic resolution details on how to improve future iterations of IPMK inhibitors.

## EXPERIMENTAL SECTION

### Protein expression and purification

Recombinant human IP6K1 and the human IPMK were prepared as previously described ^19^. The purity of these proteins was estimated to be >80% as judged by SDS-PAGE. The purified proteins were stored in aliquots at -80°C.

### IC_50_ determinations by in vitro kinase assays

For all enzyme assays, a serial dilution of the enzyme was performed in the kinase assay to determine the linear range of enzyme concentration to be used.

### IP6K kinase activity assay

An enzyme-coupled assay was used to measure IP6K activity ^19^. Enzyme activity was assayed at 37°C in 50 μL reaction mixtures containing 100nM IP6K1, 5.0 µM human Dipp1, 20 mM HEPES (pH 7.2), 100 mM KCl, 3.5 mM MgCl_2_, 20 μM EDTA, 25 μM InsP_6_ and 500 μM ATP for 60-120 min. P_i_ release was determined with a malachite green colorimetric assay. Reactions were quenched by addition of 100 μL of phosphate detection reagent (36:1 v/v of 2.6% sodium molybdate in 2.5 M HCl : 0.126% malachite green chloride). Phosphate (Pi) release was quantified from the absorbance at 620 nm using appropriate standards.

### IPMK kinase activity assay

We prepared ^33^P-1,3,4,5-IP_4_ by incubating 1,4,5-IP_3, 33_P-ATP and IPMK, isolated the product with 30 Kd MW cut-off filter. Each reaction contained trace amounts (400,000 disintegrations per minute (dpm)) of ^33^P-1,3,4,5-IP_4_ in a 100 μL incubations containing 20 mM HEPES (pH 7.2), 100 mM KCl, 3.5 mM MgCl_2_, 20 μM EDTA, 1.0 mM ATP, 1 μM 1,3,4,5-IP_4_, plus test compound in DMSO (or vehicle control) plus 2 nM IPMK. Assays were analyzed by HPLC, using a PartiSphere SAX 120 Å, 5 µm, 4.6 x 125 mm HPLC column with a 250 µL injection volume. The elution gradient was generated by mixing Buffer A (1 mm Na_2_EDTA) with Buffer B (Buffer A plus 2.5 M NH_4_H_2_PO_4_, pH 3.9) and monitored by a Beta-RAM 6 in-line scintillation detector, 1.0 mL/min elute was mixed with 0.5 or 2.5 mL/min mono-flow scintillation liquid for ^33^P or ^3^H, respectively. Radiochromatography data were collected and analyzed using Laura™ software (v6.1.2.36).

### IP3K kinase activity assay

Each reaction contained 1.0 µM [^3^H]-1,4,5-InsP_3_ (approx. 30,000 dpm; American Radiolabeled Chemicals, Inc., ART 0270) in a 100 μL incubations containing 20 mM HEPES (pH 7.2), 100 mM KCl, 3.5 mM MgCl_2_, 20 μM EDTA, 1.0 mM ATP, 1.0 μM 1,4,5-InsP_3_, plus test compound in DMSO (or vehicle control) and 0.25 nM IP3KA. Reactions were quenched after 60 min by addition of 2 volumes of 0.2 M NH_4_H_2_PO_4_, pH 3.9 and 20 mM EDTA and stored at 27°C. assays were analyzed by HPLC, using a PartiSphere SAX 120 Å, 5 µm, 4.6 x 125 mm HPLC column with a 250 µL injection volume. The elution gradient was generated by mixing Buffer A (1 mm Na_2_EDTA) with Buffer B (Buffer A plus 2.5 M NH_4_H_2_PO_4_, pH 3.9) and monitored by a Beta-RAM 6 in-line scintillation detector, 1.0 mL/min elute was mixed with 0.5 or 2.5 mL/min mono-flow scintillation liquid for ^33^P or ^3^H, respectively. Radiochromatography data were collected and analyzed using Laura™ software (v6.1.2.36).

### Isothermal titration calorimetry (ITC)

Isothermal calorimetry experiments were performed using a MicroCal PEAQ-ITC (Malvern Panalytical) with 15μM recombinant human IPMK in the sample cell and 150μM of inhibitor **1** in the syringe, each of which was maintained at 25°C in buffer containing 20mM Tris-HCl, pH 7.2, 150mM KCl, 0.05mM EDTA and 0.8mM MgCl_2_. The sample cell (volume = 204 µL) and the syringe were cleaned before each run. Thermograms were constructed from 20 injections, each of which involved 2.0 µL of ligand delivered for 4.0s, with an equilibration time of 150-300s between each injection. The stirring speed was set to 750 rpm. Data were fitted to a single binding site model using the analysis software provided by the manufacturer, as supported by the X-ray crystallography. At least three runs were performed.

### X-ray crystallography structural studies

Crystals of human apo-IPMK were prepared as described previously (14). Complex crystals were produced by soaking apo crystals into a mixture of 2–10 mM compounds with 35% (w/v) PEG 400, 0.1 M Li2SO4, 100 mM HEPES (pH 7.5) at 25°C for 3 days. Diffraction data were collected using APS beamlines 22-ID and 22-BM. All data were processed with the program HKL2000. The crystal structures were determined by using rigid body and direct Fourier synthesis, and refined with the equivalent and expanded test sets by using programs in the CCP4 package. Rotamer outliers and clash scores were calculated using Phenix. The molecular graphics representations were prepared with the program PyMol (Schrödinger, LLC). Atomic coordinates and structure factors for human IPMK/compound co-complexes have been deposited with the Protein Data Bank with the following accession codes 8V6W(**1**), 8V6X(**2**), 8V6Y(**3**), 8V6Z(**4**), 8V70(**6**), 8V71(**7**), 8V72(**8**), 8V73(**9**), 8V74(**10**), 8V75(**11**), 8V77(**12**), 8V78(**13**), 8V79(**14**), 8V76(**15**). All structural refinement data are shown in Supplementary Table S1.

## Acknowledgements

The authors acknowledge the kind gift of all compounds used in this study from Prof. Xiaodong Wang, University of North Carolina, Chapel Hill. This work was supported by the National Institute of Environmental Health Sciences (S.B.S.), an NIEHS Intramural Research Program Grant (ZIA ES 103247) to Robin E. Stanley; and a National Institutes of General Medical Sciences (NIGMS) R35 GM156389 to R.D.B.

## Abbreviations Used

dpm: disintegrations per minute
HPLC: high-performance liquid chromatography
Ins(1,4,5)P_3_: inositol 1,4,5-trisphosphate
InsP_5_: inositol pentakisphosphate
InsP_6_: inositol hexakisphosphate
5-InsP_7_: 5-diphosphoinositol pentakisphosphate
IP6K: inositol hexakisphosphate kinase
ITC: isothermal titration calorimetry
TNP: *N*2-(*m*-Trifluorobenzyl)-*N*6-(*p*-nitrobenzyl)purine

