## Supplementary Data for "Structural rationalization of IPMK inhibitor potency" (from another version of this preprint: version 1, 2025-08-16)

**Table 1, continued** Data collection and refinement statistics

| PDB Accession Codes | 8V6W | 8V6X | 8V6Y | 8V6Z | 8V70 | 8V71 | 8V72 |
|---|---|---|---|---|---|---|---|
| **Compound** | 1 | 2 | 3 | 4 | 6 | 7 | 8 |
| **Space group** | $P4_22_12$ | $P4_22_12$ | $P4_22_12$ | $P4_22_12$ | $P4_22_12$ | $P4_22_12$ | $P4_22_12$ |
| **Cell parameters a, c (Å)** | 77.7, 84.8 | 78.2, 84.3 | 78.4 85.2 | 77.9, 85.6 | 78.1, 84.8 | 78.3, 85.1 | 77.9, 85.0 |
| **Resolution (Å)*** | 1.95 (1.98) | 1.75 (1.78) | 1.7 (1.73) | 1.85 (1.88) | 1.85 (1.88) | 1.70 (1.73) | 2.0 (2.03) |
| **Rmeas*** | 0.108 (0.816) | 0.069 ( 0.693) | 0.042 (0.793) | 0.119 (0.819) | 0.092 (0.813) | 0.056 (0.783) | 0.127 (0.713) |
| **I/σ I*** | 19.3 (2.4) | 42.5 (5.0) | 65.2 (4.0) | 20.4 (2.2) | 20.9 (2.4) | 52.0 (4.8) | 20.8 (3.6) |
| **Completeness (%)*** | 99.7 (100.0) | 99.7 (99.9) | 99.3 (99.9) | 99.1 (98.6) | 99.7 (100.0) | 99.5 (100.0) | 99.9 (100.0) |
| **Redundancy*** | 10.1 (10.7) | 15.2 (14.8) | 14.2 (14.3) | 11.4 (8.6) | 7.8 (7.2) | 16.4 (16.5) | 12.5 (11.1) |
| **Refinement** | | | | | | | |
| **No. reflections*** | 19354 (1802) | 26756 (2566) | 29501 (2813) | 22642 (2061) | 22620 (2002) | 29485 (2863) | 18216 (1751) |
| **$R_{work}$*** | 0.162 (0.193) | 0.189 (0.233) | 0.144 (0.190) | 0.139 (0.231) | 0.190 (0.248) | 0.183(0.226) | 0.172 (0.208) |
| **$R_{free}$*** | 0.228 (0.271) | 0.242 (0.275) | 0.191 (0.271) | 0.229 (0.314) | 0.239 (0.269) | 0.230 (0.330) | 0.243 (0.309) |
| **No. atoms** | | | | | | | |
| **Protein** | 1941 | 1963 | 1992 | 1969 | 1958 | 1983 | 1968 |
| **ligands** | 22 | 23 | 20 | 22 | 25 | 24 | 30 |
| **solvent** | 110 | 179 | 182 | 157 | 130 | 163 | 134 |
| **R.M.S. deviations** | | | | | | | |
| **Bond length (Å)** | 0.007 | 0.007 | 0.009 | 0.006 | 0.006 | 0.007 | 0.006 |
| **Bond angle (°)** | 1.24 | 1.14 | 1.39 | 1.01 | 1.11 | 1.14 | 1.06 |
| **B-factors (Å$^2$)** | | | | | | | |
| **Protein** | 33.5 | 38.2 | 25.9 | 26.1 | 30.7 | 23.2 | 33.1 |
| **ligand** | 26.1 | 27.9 | 17.7 | 20.4 | 29.2 | 14.6 | 31.7 |
| **solvent** | 41.9 | 53.9 | 38.4 | 36.8 | 48.2 | 38.1 | 45.4 |
| **Ramachandran** | | | | | | | |
| **favored (%)** | 99.2 | 97.4 | 99.1 | 97.4 | 96.5 | 99.1 | 98.7 |
| **allowed (%)** | 0.4 | 2.6 | 0.9 | 2.6 | 2.6 | 0.9 | 1.3 |
| **outliers (%)** | 0.4 | 0.0 | 0.0 | 0.0 | 0.9 | 0.0 | 0.0 |
| **Rotamer outliers (%)** | 1.4 | 0.0 | 0.9 | 0.0 | 0.9 | 0.9 | 0.0 |
| **Clash score** | 3.1 | 4.3 | 3.5 | 1.8 | 2.8 | 2.8 | 2.5 |

*The numbers in parentheses are given for the highest-resolution shells.

**Table 1, continued** Data collection and refinement statistics

| PDB Accession Codes | 8V73 | 8V74 | 8V75 | 8V77 | 8V78 | 8V79 | 8V76 |
|---|---|---|---|---|---|---|---|
| Compounds | 9 | 10 | 11 | 12 | 13 | 14 | 15 |
| Space group | $P4_22_12$ | $P4_22_12$ | $P4_22_12$ | $P4_22_12$ | $P4_22_12$ | $P4_22_12$ | $P4_22_12$ |
| Cell parameters a, c (Å) | 77.8, 84.9 | 77.8, 85.5 | 77.8, 85.3 | 77.8, 85.3 | 78.2, 85.3 | 77.8, 85.0 | 78.5, 85.5 |
| Resolution (Å)* | 1.9(1.93) | 1.85(1.88) | 1.85(1.88) | 1.95(1.98) | 1.95(1.98) | 1.95(1.98) | 1.95(1.98) |
| Rmeas* | 0.113(0.958) | 0.110(0.912) | 0.085(0.851) | 0.110(0.864) | 0.111(0.661) | 0.094(0.816) | 0.085(0.969) |
| I/σ I* | 22.8(2.5) | 29.9(3.3) | 36.9(3.6) | 25.9(3.0) | 23.7(3.4) | 25.9(2.2) | 33.8(3.1) |
| Completeness (%)* | 99.7(100.0) | 99.9(99.6) | 99.8(99.9) | 100.0(100.0) | 99.8(99.9) | 97.3(90.9) | 98.8(98.0) |
| Redundancy* | 14.2(11.4) | 18.6(16.3) | 17.8(15.5) | 14.2(12.7) | 12.7(10.4) | 14.8(8.8) | 17.6(14.3) |
| Refinement | | | | | | | |
| No. reflections* | 20844(1851) | 22992(2215) | 22933(2213) | 19555(1821) | 19526(1893) | 18676(1435) | 19620(1885) |
| $R_{work}$* | 0.160(0.213) | 0.158(0.202) | 0.159(0.231) | 0.161(0.200) | 0.171(0.208) | 0.175(0.221) | 0.161(0.263) |
| $R_{free}$* | 0.229(0.258) | 0.217(0.266) | 0.221(0.299) | 0.217(0.303) | 0.221(0.243) | 0.241(0.291) | 0.217(0.337) |
| No. atoms | | | | | | | |
| Protein | 1953 | 1957 | 1948 | 1965 | 1965 | 1947 | 1965 |
| ligands | 27 | 26 | 31 | 27 | 29 | 30 | 27 |
| solvent | 146 | 171 | 173 | 132 | 138 | 128 | 132 |
| R.M.S. deviations | | | | | | | |
| Bond length (Å) | 0.007 | 0.006 | 0.007 | 0.006 | 0.006 | 0.007 | 0.006 |
| Bond angle (°) | 1.26 | 1.09 | 1.09 | 1.13 | 1.09 | 1.18 | 1.13 |
| B-factors (Å$^2$) | | | | | | | |
| Protein | 28.8 | 26.4 | 26.8 | 28.4 | 28.8 | 34.4 | 28.4 |
| ligand | 26.5 | 29.9 | 21.9 | 27.2 | 27.1 | 26.7 | 27.2 |
| solvent | 35.0 | 35.4 | 38.6 | 40.7 | 42.8 | 44.5 | 40.7 |
| Ramachandran | | | | | | | |
| favored (%) | 98.7 | 97.9 | 99.1 | 98.3 | 99.1 | 97.9 | 98.3 |
| allowed (%) | 0.9 | 2.1 | 0.9 | 1.7 | 0.9 | 2.1 | 1.7 |
| outliers (%) | 0.4 | 0.0 | 0.0 | 0.0 | 0.0 | 0.0 | 0.0 |
| Rotamer outliers (%) | 0.9 | 0.5 | 0.0 | 0.0 | 0.0 | 0.0 | 0.0 |
| Clash score | 3.9 | 2.1 | 2.6 | 2.6 | 2.8 | 2.8 | 2.6 |

*The numbers in parentheses are given for the highest-resolution shells.
